# Monozygotic twins discordant for schizophrenia differ in maturation and synaptic transmission

**DOI:** 10.1101/2022.05.13.491776

**Authors:** Shani Stern, Lei Zhang, Meiyan Wang, Rebecca Wright, Diogo Cordeiro, David Peles, Yuqing Hang, Ana P. D. Mendes, Tithi Baul, Julien Roth, Shashank Coorapati, Marco Boks, Hilleke Hulshoff Pol, Kristen J. Brennand, Janos M Réthelyi, René S. Kahn, Maria C. Marchetto, Fred H. Gage

**Affiliations:** Laboratory of Genetics, The Salk Institute for Biological Studies, La Jolla, CA, USA; Sagol Department of Neurobiology, Faculty of Natural Sciences, University of Haifa, Haifa, Israel; Department of Psychiatry, University Medical Center Utrecht Brain Center, Utrecht University, Heidelberglaan 100, 3584CX Utrecht, The Netherlands; Department of Experimental Psychology, Utrecht University, Heidelberglaan 1, 3584CS Utrecht, The Netherlands; Nash Family Department of Neuroscience, Friedman Brain Institute, Pamela Sklar Division of Psychiatric Genomics, Black Family Stem Cell Institute, Icahn School of Medicine at Mount Sinai, New York, NY 10029, USA; Department of Psychiatry, Department of Genetics, Yale Stem Cell Center, Yale University School of Medicine, New Haven, CT 06511, USA; Molecular Psychiatry Research Group and Department of Psychiatry and Psychotherapy, Semmelweis University, Budapest, Hungary; Department of Psychiatry, Icahn School of Medicine at Mount Sinai, New York, NY, USA. Mental Illness Research, Education and Clinical Center, James J Peters VA Medical Center, New York, NY, USA; Department of Anthropology, University of California San Diego, 9500 Gilman Drive, La Jolla, CA 92093, USA

## Abstract

Schizophrenia affects approximately 1% of the world population. Genetics, epigenetics, and environmental factors are known to play a role in this psychiatric disorder. While there is a high concordance in monozygotic twins, about half of twin pairs are discordant for schizophrenia. We characterized human-induced pluripotent stem cell (iPSC)-derived hippocampal neurons from two pairs of monozygotic twins that are discordant for a schizophrenia diagnosis. We compared the affected and the non-affected siblings and compared all of them to twin sets where none of the siblings suffered from schizophrenia. We found that the neurons derived from the schizophrenia patients were less arborized, were hypoexcitable with immature spike features, and exhibited a significant reduction in synaptic activity with dysregulation in synapse-related genes. Interestingly, the neurons derived from the co-twin siblings who did not have schizophrenia formed another distinct group that was different from the neurons in the group of the affected twin siblings but also different from the neurons in the group of the control twins. The neurons in the unaffected co-twin group were also less arborized than the neurons from controls but more arborized than those from affected siblings. Some of their spike features were immature (but less immature than neurons derived from the affected siblings). Importantly, their synaptic activity was not affected. Since schizophrenia is a genetically complex disorder, our twin study allows the measurement of neuronal phenotypes with a similar genetic background. The differences between the siblings may arise due to changes that occurred after the split of the egg into twins. Therefore, our study confirms that dysregulation of synaptic pathways, as well as changes in the rate of synaptic events, distinguishes between individuals affected with schizophrenia and unaffected individuals, even in those having a very similar genetic background.

## Introduction

The prevalence of schizophrenia is approximately 1% worldwide^1^, and full recovery of these patients is limited and the prognosis is guarded^2, 3^. Schizophrenia is defined in the *Diagnostic and Statistical Manual of Mental Disorders, Fifth Edition*, (*DSM-5*) by at least 2 of the following symptoms: delusions, hallucinations, disorganized speech, negative symptoms, and disorganized or catatonic behavior. At least 1 of the symptoms must be the presence of disorganized speech, delusions, or hallucinations. These signs and symptoms of the disturbance must persist for at least 6 months; during this time, the patient must experience at least 1 month of active symptoms and social or occupational deterioration problems over a significant period. These problems are unique and not related to other conditions, e.g. psychoactive substances or neurologic conditions.

Although many studies have been conducted to elucidate the genes and molecules related to schizophrenia, it remains unclear as to which of the candidate genes are most critical to the disease. Post-mortem studies and neuroimaging technologies have shown differences between the brains of schizophrenia patients compared to healthy individuals. A decrease in brain volume in the medial-temporal areas, changes in the hippocampus and larger ventricles and white matter tracts, and structural brain abnormalities are some of these alterations^4-6^.

Neurotransmitter abnormalities have also been elucidated, especially in the dopaminergic system, although some studies also implicate other neurotransmitter systems such as gamma-aminobutyric acid (GABA), norepinephrine, serotonin, and NMDA glutamate receptors^7-10^. In addition to a genetic contribution, the environment also plays a key role in disease etiology ^11^. This disorder is hereditary and the concordance of schizophrenia in monozygotic twins ranges from 41%-79% ^12-14^. Genome-wide association studies (GWAS) have been performed in SCZ with an approach to better understand the etiology of the disease. Many genes have been associated with schizophrenia^15-18^. Some of the associated genes include *CACNA1C, DGCR8, DRD2, MIR137, NOS1AP, NRXN1*. More than 100 SCZ-related loci have been reported, although the GWAS loci identified failed to identify the high-risk genes (HRGs) related to SCZ ^17, 19^. Single-nucleotide polymorphisms (SNPs) and their link with associated genes are often difficult to interpret, especially when the SNPs are in noncoding regions. Many methods have been attempted to identify risk genes linked to SCZ regulated by GWAS loci. For instance, integrating position weight matrix (PWM) and functional genomics ^20, 21^or topologically associated domains that are generated by chromatin interaction experiments^22^. Using large databases of GWAS, transcriptome-wide associated studies (TWAS), large web-based platforms, multi-omics data, and gene expression in statistical models were used to predict HRGs in SCZ in attempt to explain the high hereditability of SCZ and provide mechanistic insights of the disease. Expression patterns of genes in SCZ neurons may further elucidate the molecular mechanisms of the disease and drug discovery ^23^. The Psychiatry Genome Consortium wave 3 (PGC3) meta-analysis has reported 287 genomic loci containing common alleles associated with schizophrenia, highlighting the complex genetic architecture enriched by neurodevelopmental genes as well as synapse-related pathways as being of central importance ^24^.

For the past decade, human stem cell technology has been used to generate virtually any human cell type^25^, and induced pluripotent stem cell (iPSC)-based models have advanced the study of neuropsychiatry disorders such as schizophrenia, depression, autism spectrum disorder, epilepsy, bipolar disorder, and Alzheimer’s disease among others diseases^18, 26-29^. The first few studies demonstrating the generation of iPSCs from patients with schizophrenia were published a decade ago^26, 30, 31^. Some of the schizophrenia cases in these studies were described as idiopathic and others had demonstrated familial inheritance. There were no intrinsic deficits in iPSC pluripotency or self-renewal when derived from schizophrenia patients compared to controls that were reported. This finding is consistent with the fact that schizophrenia does not have large effects on embryogenesis or affect multiple organ systems. Also, there were no differences found between patient and control iPSCs in terms of induction, quantity, or morphology.

Chiang et al ^30^ were the first to publish the generation of human iPSCs from schizophrenia patients with a mutation in the DISC1 gene that is known to be linked to schizophrenia ^32^. Neuronal changes have been reported also by Brennand et al.^26^, who observed differences in iPSCs patient-derived neurons such as neuronal connectivity, decreased neurite number, and postsynaptic density protein 95(PSD95) levels that were also reduced. Gene expression profiles were also different in schizophrenia neurons with alterations in genes associated with cAMP, WNT pathways, and glutamate receptor expression. A few years later, the same lab reported that neurons that were derived from schizophrenia patients were hypoexcitable^33^. Pedrosa et al.^31^ also demonstrated the potential use of iPSCs in modeling schizophrenia, showing that derived neurons expressed chromatin remodeling proteins, transcriptional factors, and synaptic proteins that were relevant to schizophrenia. A significant delay in the reduction of endogenous *OCT4* and *NANOG* expression was shown during differentiation in the schizophrenia lines. Na^+^ channel function and GABA-ergic neurotransmission have recently been shown to be altered in neurons derived from iPSCs of schizophrenia patients^34^. These new approaches to neuropsychiatry disorders have the potential to address the problem of brain inaccessibility by providing the investigators with specific neurons of patients with proper diagnostic validity (or simply proper diagnoses).

In this study, iPSCs from monozygotic twins discordant for schizophrenia and a control group of healthy twins were differentiated into neural progenitor cells (NPCs), immature, and mature neurons. The advantage of this system is the similar genetic background of the affected and unaffected siblings, which minimizes the genetic diversity and allows us to focus on phenotypes that are specific to schizophrenia. The differentiated neurons were analyzed over a period from four weeks to two months, and the transcriptome (RNAseq), morphology, histology, and also functional assays with electrophysiology and microelectrode array (MEA) were used to compare the affected patients and their unaffected twin siblings to the control group. We found an immature state of dentate gyrus (DG) granule neurons that were derived from the schizophrenia patients, with an intermediate state in the neurons that were derived from the unaffected twins. An immature dentate gyrus was previously shown to manifest an immature molecular profile in schizophrenia subjects, as well as in various animal models^35^. Our results further support this phenotype in a human cellular model. The most pronounced phenotype was an impairment of the synaptic transmission in the schizophrenia patients measured both with electrophysiology and at the gene expression levels.

## Materials and Methods

### Patient selection

The recruitment of the subjects and biopsies were carried out in Utrecht, Netherlands at the Department of Psychiatry, UMC Utrecht. Monozygotic twin pairs discordant for schizophrenia and control monozygotic twin pairs from the U-TWIN cohort ^36^ were asked to participate in the study. Diagnosis of schizophrenia was made according to DSM-IV. Control twins were excluded if they ever met the criteria for a psychotic or manic disorder substance dependence, had a first-degree relative with schizophrenia, or if were diagnosed with a neurological disorder. All participants gave their written informed consent. The Medical Ethical Committee of the UMC Utrecht approved this study and the experiments were under the Declaration of Helsinki. IRB approval was obtained by Salk Institute and the samples were de-identified. Altogether, fibroblast cells from 5 twin pairs have been successfully reprogrammed by the staff of the Salk Institute Stem Cell Core Laboratory (subject age: 32-50 years). Two pairs of subjects are discordant for SCZD (age at onset of SZCD: 22-35 years) and 3 are healthy control twins (Figure 1).

**Figure 1.**
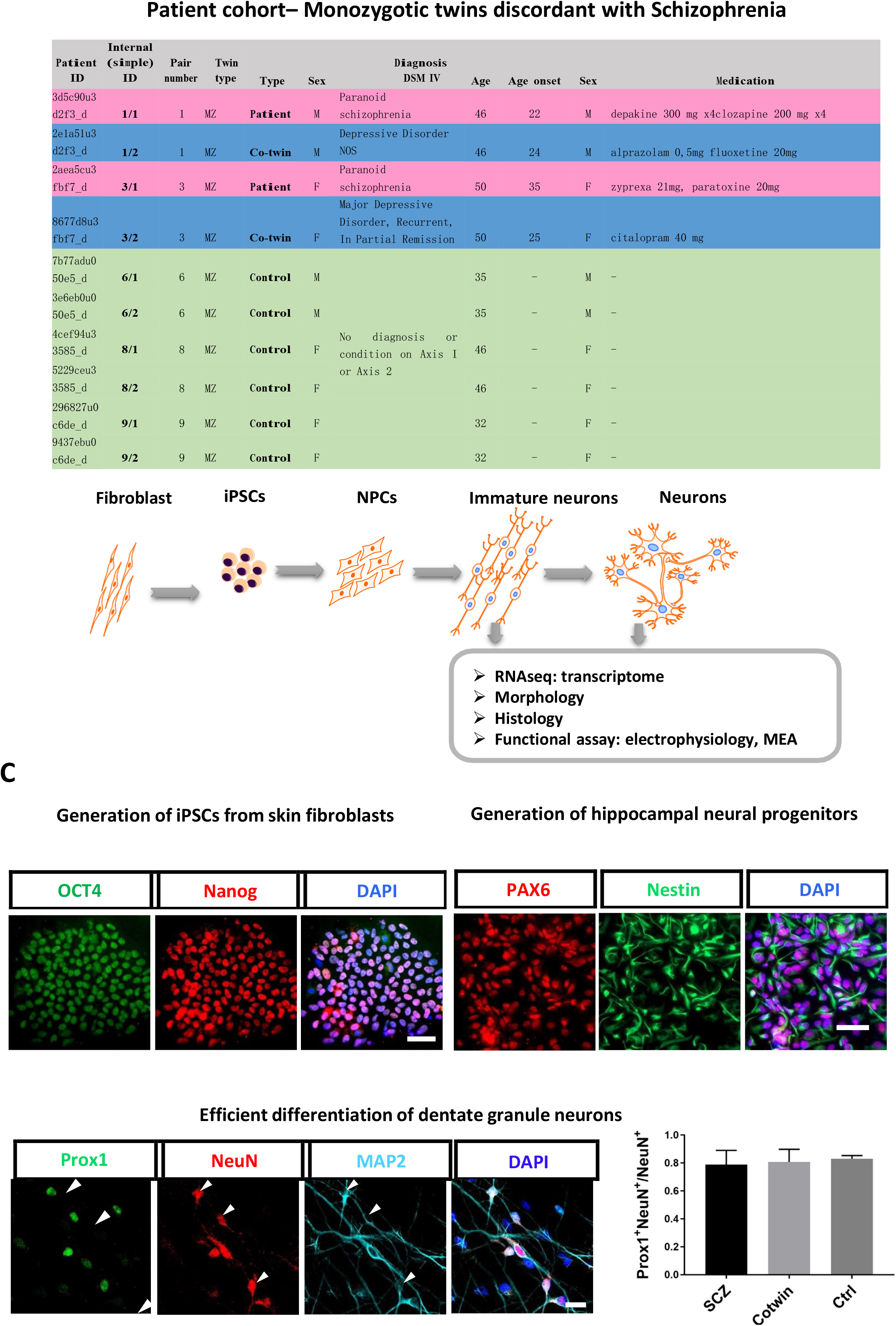
Generation of neurons from healthy twin sets and from twin sets that are discordant for schizophrenia. A. A table describing the cohort of patients and control individuals. The lines with the information of the affected patients are highlighted in pink and the unaffected twin siblings are highlighted in light blue. The green lines contain information about the healthy control twin sets. B. A schematic of the reprogramming and differentiation and the experiments performed. C. Upper row, Left – Pluripotency markers were expressed in the reprogrammed iPSCs. Upper row, Right – Neural Progenitor cells (NPCs) expressed Nestin and PAX6–NPC-specific markers. Lower row – differentiated neurons expressed MAP2 and approximately 80% expressed Prox1, a marker that is specific for the DG granule neurons in neurons derived from the 3 groups of affected twin siblings, unaffected co-twin siblings, and healthy twin sets. Scale bars are 50 μm.

### iPSC reprogramming and neuron differentiation

iPSCs were derived from fibroblasts for control and schizophrenia patients using the Cyto-Tune Sendai reprogramming kit (Invitrogen) according to the manufacturer’s instructions. iPSCs were characterized for a normal karyotype as well as for pluripotency markers as previously described^37^. iPSC colonies were cultured on Matrigel-coated plates using mTeSR medium (StemCell Technologies). Embryoid bodies (EBs) were formed by mechanical dissociation of iPSC colonies using dispase and then plating in ultra low-attachment plates. The mTeSR was replaced on the next day with DMEM/F12 (Invitrogen) supplemented with N2 and B27. For EB differentiation, floating EBs were treated with DKK1 (0.5 μg /ml), SB431542 (10 μM), noggin (0.5 μg/ml) and cyclopamine (1 μm) for 20 days. To obtain neural progenitor cells (NPCs), the embryoid bodies were plated onto poly-L-ornithine/laminin-coated plates. The rosettes were manually collected and dissociated with accutase after 1 week and plated onto poly-L-ornithine/laminin-coated plates in NPC media containing DMEM/F12, N2, B27, 1 μg/ml laminin and 20 ng/ml FGF2. To obtain mature neurons, NPCs were differentiated in DMEM/F12 supplemented with N2, B27, 20 ng/ml BDNF (Peprotech), 1 mM dibutyrl-cyclicAMP (Sigma), 200 nM ascorbic acid (Sigma), 1 μg/ml laminin and 620 ng/ml Wnt3a (R&D) for 3 weeks. Wnt3a was removed after 3 weeks. Neurons were infected with the Prox1∷eGFP lentiviral vector at 8 days of differentiation and the experiments were performed in the Prox1-positive neurons. The neuronal cultures were dissociated and replated on poly-L-ornithine/laminin coated coverslips at 2 weeks using accutase.

### Immunohistochemistry

Cells on glass coverslips were fixed in 4% paraformaldehyde for 15 min. The cells were then blocked and permeabilized in PBS containing 0.1–0.2% Triton X-100 and 10% horse serum. Coverslips were then incubated with the primary antibody in the blocking solution overnight at 4 °C. The coverslips were washed in Tris-buffered saline and incubated with the secondary antibodies for 30 min at room temperature and counterstained with DAPI. The coverslips were then washed and mounted on slides using Fluoromount-G (Southern Biotech), and dried overnight in the dark. The antibodies used for pluripotency and neuronal characterization were anti-hOct4 (Cell Signaling, Danvers, MA, USA 2840S), anti-hSox2 (Cell Signaling 3579S), anti-hNanog antibody (Cell Signaling 4903S), Tra-1-81 (Cell Signaling; cat. no. 4745S), anti-MAP2 (Abcam 5392) and anti-GFP (Abcam, San Francisco, CA, USA; cat. no. 6673), PAX6 (Abcam ab109233), Nestin (Abcam ab105389), NeuN(Abcam ab177487), and PROX1 (Millipore MAB5654).

### Sholl analysis

Differentiated neurons were traced using Neurolucida (MBF Bioscience, Williston, VT). Only neurons that were PROX1-positive were included in this analysis. The morphology of the neurons was quantified using Neurolucida Explorer (MBF Bioscience, Williston, VT). Sholl analysis was performed using Neurolucida Explorer’s sholl analysis option. This analysis specified a center point within the soma and created a grid of concentric rings around it with radii increasing in 10-µm increments. Neuronal complexity was determined by recording the number of intersections within each ring

### Electrophysiology

The coverslips with neuronal cultures were transferred to a recording chamber at 2 time points - 4 weeks and 2 months - with artificial cerebrospinal fluid (ACSF) containing (in mM): 10 HEPES, 4 KCl, 2 CaCl2,1 MgCl2, 139 NaCl, 10 D-glucose (310 mOsm, pH 7.4). Whole-cell patch-clamp recordings were performed from Prox-positive neurons. Patch electrodes were filled with an internal solution containing (in mM): 130 K-gluconate, 6 KCl, 4 NaCl, 10 Na-HEPES, 0.2 K-EGTA, 0.3 GTP, 2 MgATP, 0.2 cAMP, 10 D-glucose, 0.15% biocytin and 0.06% rhodamine. The pH and osmolarity of the internal solution were adjusted to a pH of 7.4 and an osmolarity of 290 mOsm. The signals were amplified with a Multiclamp700B amplifier (Sunnyvale, CA, USA) and recorded with Clampex 10.3 software (Axon Instruments, Union City, CA, USA). Data were acquired at a sampling rate of 20 kHz and analyzed using Clampfit-10 and the software package MATLAB (release 2014b; The MathWorks, Natick, MA, USA). All measurements were conducted at room temperature.

### Analysis electrophysiology

#### Total evoked action potentials

The cells were typically held in current-clamp mode at −60 mV and current injections were given starting 12 pA below the holding current, in 3-pA steps of 400 ms in duration. A total of 20 depolarization steps were injected. Neurons with a holding current of more than 50 pA were discarded from the analysis. The total number of action potentials was counted in 20 depolarization steps.

#### Action potential shape analysis

The first evoked action potential was used for this analysis (with the minimal injected current needed for an action potential to occur). The spike threshold was the membrane potential of the first maximum in the second derivative of the voltage by time. The fast afterhyperpolarization (AHP) amplitude was calculated as the difference between the threshold for an action potential and the membrane potential 5 ms after the membrane potential returned to cross the threshold value after the action potential resumed. The spike amplitude was calculated as the difference between the maximum membrane potential during an action potential and the threshold. The spike width was calculated as the time it took the membrane potential to reach half the spike amplitude in the rising part of the spike to the descending part of the spike (full-width at half-maximum).

#### The membrane resistance

was calculated around the resting membrane potential by measuring the current at −70 mV and then at −50 mV. The membrane resistance was calculated by dividing 20 mV by the difference in these currents.

#### Sodium and potassium currents

The sodium and potassium currents were acquired in voltage-clamp mode. The cells were held at −60 mV, and voltage steps of 400 ms were then given in the range of −90 to 80 mV. The fast potassium current measurement was obtained as the maximal current within a few milliseconds after the depolarization step. The slow potassium currents were obtained at the end of the 400 ms depolarization step. The sodium current was obtained by subtracting the minimum current, representing the inward sodium current from the current after stabilizing from the transient sodium current.

#### Excitatory postsynaptic currents recordings

Excitatory postsynaptic currents (EPSCs) were measured by voltage clamping the cells at -60 mV after application of 40 μM of bicuculline, a GABA_A_ antagonist. The analysis was performed using semi-manual Matlab scripts.

### Multiwell microelectrode array (MEA) recordings and analysis

To record the spontaneous activity of neurons derived from schizophrenia twins and healthy twins, neurons at 31-day differentiation were seeded in 96 wells MEA plates from Axion Biosystems (San Francisco, CA, USA). Each subject’s cells were plated in replicates of 6 and seeded with 10,000 neurons each well. Cells were fed every 2-3 days and electrical activity was recorded every 3-4 days from day 37 of neuronal differentiation using the maestro MEA system and Axis software (Axion Biosystems). Voltages were recorded at a frequency of 12.5 kHz and bandpass filtered between 10 Hz and 2.5 kHz. Spike detection was performed using an adaptive threshold set to 5.5 standard deviations above the mean activity of each electrode. Following 5 minutes of plate adaptation time, recordings were performed for 10 minutes. Multielectrode data analysis was performed using the Axion Biosystems Neural Metrics Tool, which calculated standard spike-related measurements. Bursts were detected with an adaptive Poisson algorithm for high spiking activity that occurred on a single electrode. Variables were averaged across subject replicates and plotted by groups for each day of the recordings. A two-way ANOVA was used to compare of the electrical activity in the 3 groups.

### FACS sorting

We have prepared the neurons for RNA-seq according to a published FACS protocol ^38^. Briefly, five-week-old neurons were dissociated and CD184−/CD44−/CD15-/CD24+ cells (Miltenyi Biotec, cat. number 130-103-868, 130-113-334, 560828, 130-099-399, respectively) were collected and subjected to Stranded mRNA(PolyA+)-Seq Library Prep. A total of 1000 ng of RNA was used for library preparation using the Illumina TruSeq RNA Sample Preparation Kit. The libraries were sequenced on Illumina HiSeq2500 with 50 bp single-end reads.

### RNA sequencing analysis

The raw fastq reads underwent sequence alignment using the Spliced Transcripts Alignment to a Reference (STAR)^39^ and HOMER (http://homer.ucsd.edu/homer/ngs/rnaseq/index.html). Both raw count and tpm quantified matrix for all experiments were generated. PCA and heatmap were generated based on the whole tpm matrix in R 3.6.1 using gplots, and DEG analysis was performed using DESeq2 ^40^. Pooled schizophrenia (SZ) patients, as well as non-affected co-twins, were compared with control twin samples. Also, SZ patients were compared with their corresponding co-twin samples respectively. Heatmap of the DEG genes were generated in R, and Functional Enrichment for the DEG genes was performed on WebGestalt (http://www.webgestalt.org/). Also, the Venn diagrams were generated on the online Venn webtool (https://bioinformatics.psb.ugent.be/webtools/Venn/).

### GWAS Genes and DEGs intersection

We used GWAS Catalogue (https://www.ebi.ac.uk/gwas/) and searched for the term “schizophrenia”. We performed an intersection between the GWAS genes and the DEGs genes. The intersection genes were computed by comparing the symbols of the GWAS genes and DEGs genes (after the symbol standardization). The intersected genes graph was generated by Matlab’s wordcloud function, down-regulated genes were painted in blue, and up-regulated genes in red. The font size is proportional to the number of GWAS studies that included this gene with p-value < 0.01.

### Statistical analysis

The default analysis when not specified was the student t-test. We used ANOVA tests in several measurements and specified this in the Results section.

## Results

### All patients, as well as control lines, differentiate with high efficiency into hippocampal DG granule neurons

Fibroblasts were produced after a skin biopsy from a total of 2 pairs of monozygotic twins discordant for schizophrenia and 3 pairs of control (healthy) twins. The data were partitioned into 3 groups: the affected twin (2 patients), the unaffected twin sibling (2 unaffected co-twins), and 6 control individuals from the healthy twins. Figure 1A presents the details about the patients, their unaffected co-twin, and the control twins. Sendai reprogramming was performed on the fibroblasts and the patients, their co-twin, and the control twins and iPSCs were prepared. The iPSCs from all the lines exhibited pluripotency markers (Fig. 1C, left). We next prepared hippocampal NPCs^27^ and, from these, we differentiated neurons and performed experiments on these neurons at 2 maturation time points (Fig. 1B for the schematics). Figure 1C, right, presents immunocytochemistry performed on the NPCs for specific markers. All the lines expressed NPC-specific markers. Figure 1C (bottom row) presents the efficient generation of DG granule neurons with approximately 80% Prox1 positive neurons.

### DG neurons derived from schizophrenia patients are less arborized and exhibit delayed maturation, while neurons derived from the co-twin exhibit a mid-state between the other 2 groups

We next performed imaging of the DG neurons derived from the 3 groups (affected twins, unaffected co-twins, and controls) of rhodamine-filled neurons at 8 weeks post-differentiation. The neurons were filled with rhodamine during patch-clamp recordings. Sholl analysis and capacitance measurements reveal that the neurons derived from the schizophrenia-affected twin siblings were the least arborized of all the 3 groups. Fig. 2A presents example traces of rhodamine-filled neurons. Figure 2B presents the average sholl analysis of n=16 control neurons, n=26 co-twin neurons, and n=29 affected neurons. Figure 2C presents the number of branches in each of the groups which were significantly smaller in the schizophrenia neurons compared to the neurons derived from the co-twins. Figure 2D presents the maximal branch length, which was significantly smaller in the schizophrenia neurons compared to the neurons derived from the co-twins, and borderline significant when compared to the control neurons (p=0.054). We have also analyzed the capacitance of the membrane of the neurons that were measured during the electrophysiology experiments from the 3 groups when the neurons were at 8 weeks after the start of the differentiation. The neurons derived from the schizophrenia patients had the smallest average capacitance, whereas the neurons derived from the unaffected co-twin had an average capacitance that was in between the affected twin and the control average capacitances (Figure 2E-F). Since the membrane capacitance was proportional to the surface area of the neuron, this measurement further supported the finding that the neurons derived from the schizophrenia patients were smaller and less arborized, and the neurons derived from the unaffected co-twin were in between the other two groups.

**Figure 2.**
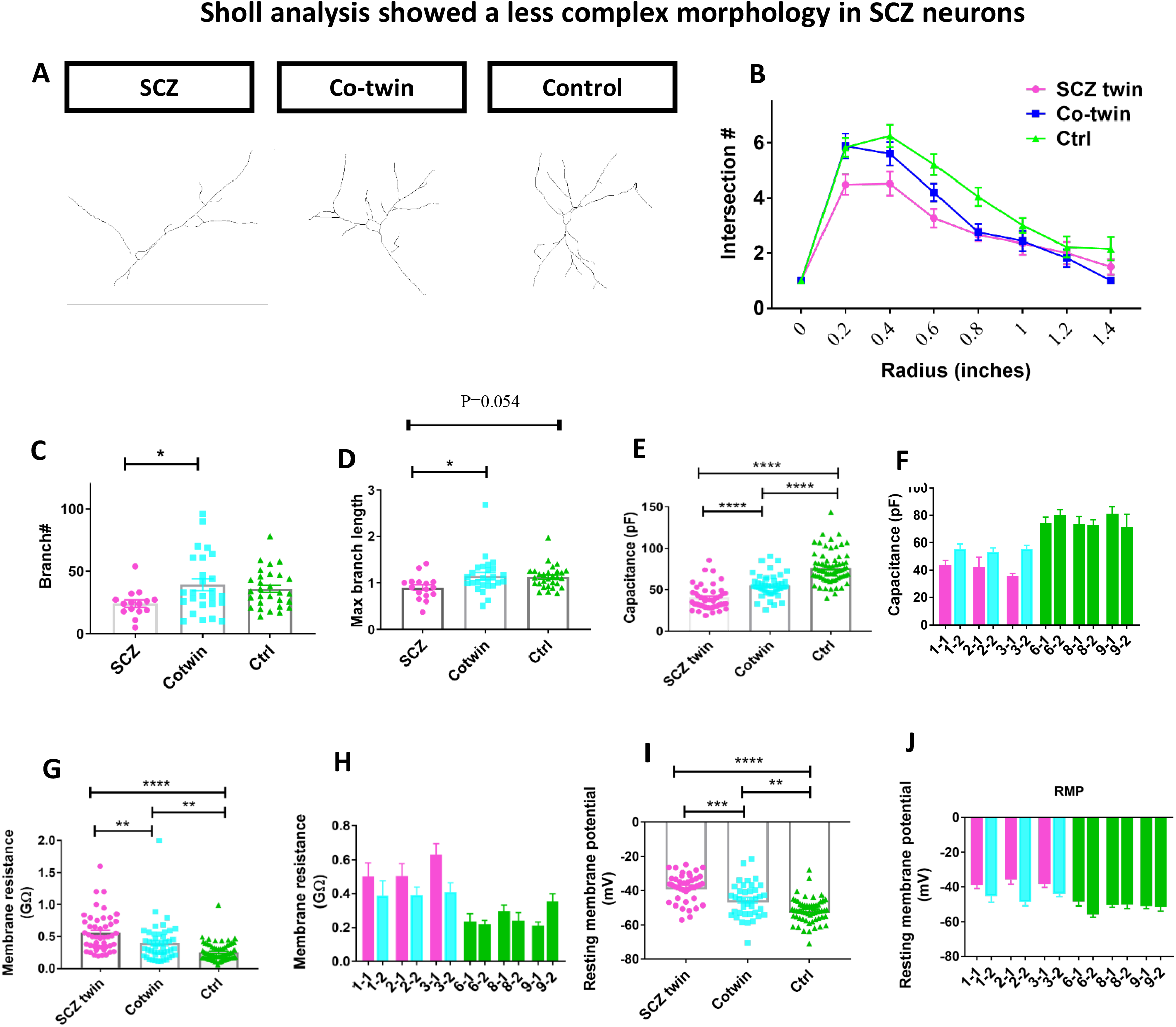
Immature features of neurons derived from the schizophrenia patients and an intermediate state of the neurons derived from the unaffected co-twin siblings. A. Example traces of a schizophrenia, co-twin, and a control neuron. B. Sholl analysis shows that neurons derived from the schizophrenia patients are less arborized than the neurons derived from their unaffected twin siblings, but both groups are less arborized than neurons derived from the healthy twin sets. C. The number of branches was reduced in the schizophrenia neuorns compared to the neurons derived from their co-twin. D. The maximum number of branches was reduced in the schizophrenia neurons compared to the neurons derived from the co-twins and borderline significant compared to the control neurons. E-F. The neurons derived from the affected twin siblings had the smallest capacitance. The neurons derived from the unaffected twin siblings had a higher capacitance than the affected twin siblings, but smaller than the healthy control twins. G-H. The neurons derived from the affected twin siblings had the highest membrane resistance. The neurons derived from the unaffected co-twin siblings had a smaller membrane resistance than the affected twin siblings but larger than the healthy control twins. I-J. The neurons derived from the affected twin siblings had the most depolarized resting membrane potential. The neurons derived from the unaffected co-twin siblings had a resting membrane potential that was less depolarized than the affected twin siblings but more depolarized compared to the healthy control twins. ** represents p<0.01, *** represents p<0.001, **** represents p<0.0001.

The membrane resistance also signifies the maturity of the neurons and is directly affected by ion channels such as the inward rectifying potassium channels and other ion channels that are open at the resting membrane potential. Measurements of the membrane resistance also revealed that the 3 groups (affected, unaffected co-twin, and control) were different from one another (Figure 2G-H). The affected twin neurons had the highest resistance (0.56±0.07 GΩ), whereas the unaffected co-twin group had an average membrane resistance that was in between the affected neurons and the control neurons (0.46±0.06 GΩ). The control neurons had the lowest average membrane resistance (0.24±0.01 GΩ). The high membrane resistance of the neurons derived from the schizophrenia patients means that they had the least number of ion channels that were open at the resting membrane potential such as the inward rectifying ion channels and Hyperpolarization-activated cyclic nucleotide-gated (HCN) channels. This finding further supports the observation that such neurons were delayed in their maturation. The resting membrane potential is another measure of neuronal maturation and is also related to the number of ion channels on the membrane. In this measure, too, we found that the groups were different from one another (Figure 2I-J). The affected twin group had the most depolarized resting membrane potential (−40.6±1.8 mV), whereas the unaffected twin group was again in the middle among the other 2 groups (−46.3±1.7 mV), and the control twins had the least depolarized membrane potential (−52.8±0.8 mV), indicating the control group displays the most mature state.

### Transcriptional changes in DG neurons derived from the schizophrenia patients compared to their unaffected co-twin and compared to the control twins

To further understand the mechanisms involved in DG neuronal changes in schizophrenia, we next performed RNA sequencing on RNA that was extracted from DG granule neurons sorted based on GFP (see methods). The neurons were dissociated and RNA was prepared approximately 7-8 weeks post-differentiation. For the first analysis, neurons derived from the affected twins (the affected and unaffected co-twins) were pooled together and compared to neurons derived from the healthy twin pairs. Figure 3A presents a PCA on the left, showing some separation between the affected twin pairs and the non-affected twin pairs. On the right, a heatmap of the top 100 differentially expressed genes is presented. Running Gene Ontology (GO), we found a few dysregulated pathways that are presented in Figure 3B. These include DG development and anterograde trans-synaptic signaling. These dysregulated pathways give further support to our morphological and electrophysiological findings. The canonical wnt pathway is also dysregulated, providing more evidence linking this pathway to neuropsychiatric disorders ^41-44^. When we compared the twin affected by schizophrenia to the unaffected co-twin (2 comparisons), we also found a few hundred differentially expressed genes. Figure 3C (i-ii) presents the heatmaps of the top 100 differentially expressed genes in 2 of the discordant twin pair. Figure 3C (iii-iv) show common genes that are differentially expressed in our dataset and appear in published GWAS. Down-regulated genes are shown in blue, and up-regulated genes in red. The most pronounced gene that is dysregulated in both pair of twins and also comes up as associated in several GWAS is the NRGN gene. Interestingly, this gene encodes a postsynaptic protein kinase substrate that binds to calmodulin in the absence of calcium. Figure 3D presents the shared differentially expressed genes between control and schizophrenia-affected patients. The 3 different comparisons (2 affected twin pairs vs. healthy twin pairs, and the 2 comparisons of the affected vs. unaffected twins discordant for schizophrenia) showed few shared genes.

**Figure 3.**
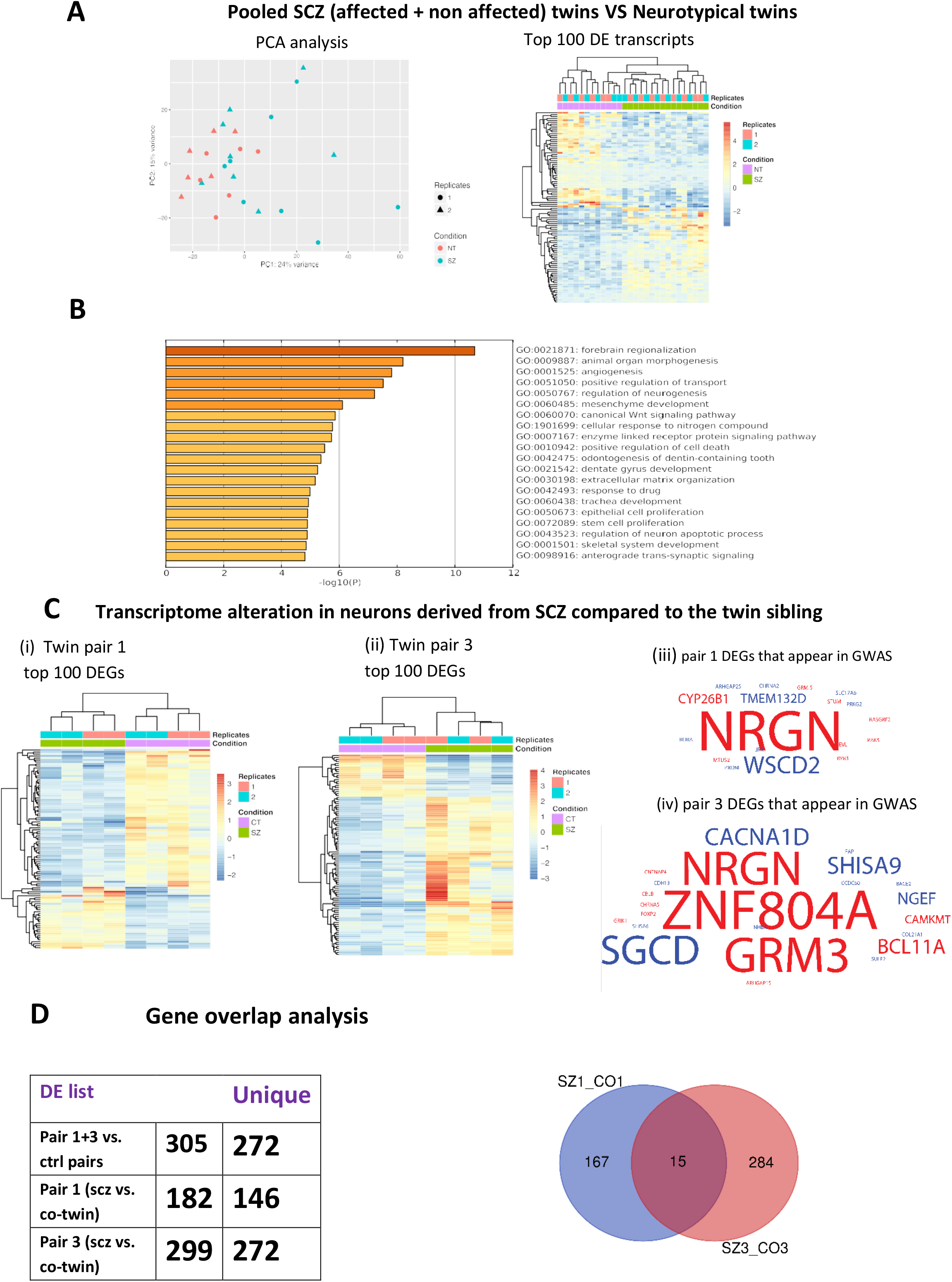
Dysregulated pathways between the affected twin set and the control twin sets include synapse-related pathways, Wnt signaling pathways, and DG development pathways. A. A total of 829 differentially expressed genes were found between the affected twin sets (both twins) and the healthy twins. A PCA plot of the gene expression data shows a partial separation into 2 groups of neurotypical and schizophrenia (affected and unaffected co-twins data pooled together) neurons. On the right, a heatmap of the top 100 dysregulated genes is presented. B. The top dysregulated pathways between the neurons derived from the affected twin sets (both the siblings) compared to the neurons derived from the healthy twin sets are presented. C. (i-ii) Heatmaps of the differentially expressed genes in the two affected twin sets (a comparison between the affected and unaffected twin) (iii-iv) Genes that appear as DEGs in our dataset as well as in GWAS. Down-regulated genes are presented in blue, and up-regulated genes in red. The font size is proportional to the number of GWAS studies that included this gene with p-value < 0.01. D. Three different comparisons were performed. The affected twins (both siblings) compared to the healthy twins, the affected twin vs. their unaffected co-twin sibling in twin set 1, and similarly in twin set 3. Most of the comparisons reveal unique genes. However, pathways that are related to synaptic signaling are affected in all 3 comparisons.

### Reduced rate of excitatory postsynaptic currents in neurons derived from the affected twins

We voltage-clamped the neurons at -60 mV to measure spontaneous excitatory postsynaptic currents (sEPSCs) after the application of bicuculine to block inhibitory postsynaptic currents. The neurons derived from the patients with schizophrenia had a drastic reduction in the EPSC frequency when compared to their unaffected co-twins or the healthy twin pairs (Fig. 4A left for representative traces and Fig. 4A right for averages over 23 neurons derived from the schizophrenia patients, 20 neurons derived from the unaffected co-twin, and 32 neurons derived from the healthy twins). The average amplitude of EPSCs was not changed between neurons derived from the 3 groups of control, unaffected co-twins, and affected twins with schizophrenia (Fig. 4A right). The data indicate a pre-synaptic deficit in schizophrenia neurons, which is consistent with previous reports^30^. Going back to the transcriptomics analysis, we next plotted the top 20 differentially expressed genes between the neurons derived from the affected twins (both affected and unaffected twin siblings) (Figure 4B, left). Of these 20 genes, 6 were synapse-related (marked in purple). The HTR7 gene, a serotonin receptor, marked in red, is associated with schizophrenia^45, 46^. We also performed GO analysis on the comparisons between neurons derived from each affected twin and his unaffected co-twin. The dysregulated pathways were highly shared when comparing the affected and unaffected twin sets. For pair number 1, the 3 dysregulated pathways were “Synapse” (p=0.000083, FDR=0.002), “Cell junction” (p=0.00014, FDR=0.002), and “Postsynaptic cell membrane” (p=0.00017, FDR=0.002). In pair 3, the most dysregulated functional annotations were similarly “Cell junction” (p=0.0072, FDR=0.17), “Postsynaptic cell membrane” (p=0.0096, FDR=0.17), and “Synapse” (p=0.021, FDR=0.25). Overall, our results demonstrate how synaptic impairment is highly involved in schizophrenia. The changes that we see in the electrophysiology relate more to the pre-synapse, but compensation mechanisms may relay these changes to the post-synapse ^47^.

**Figure 4.**
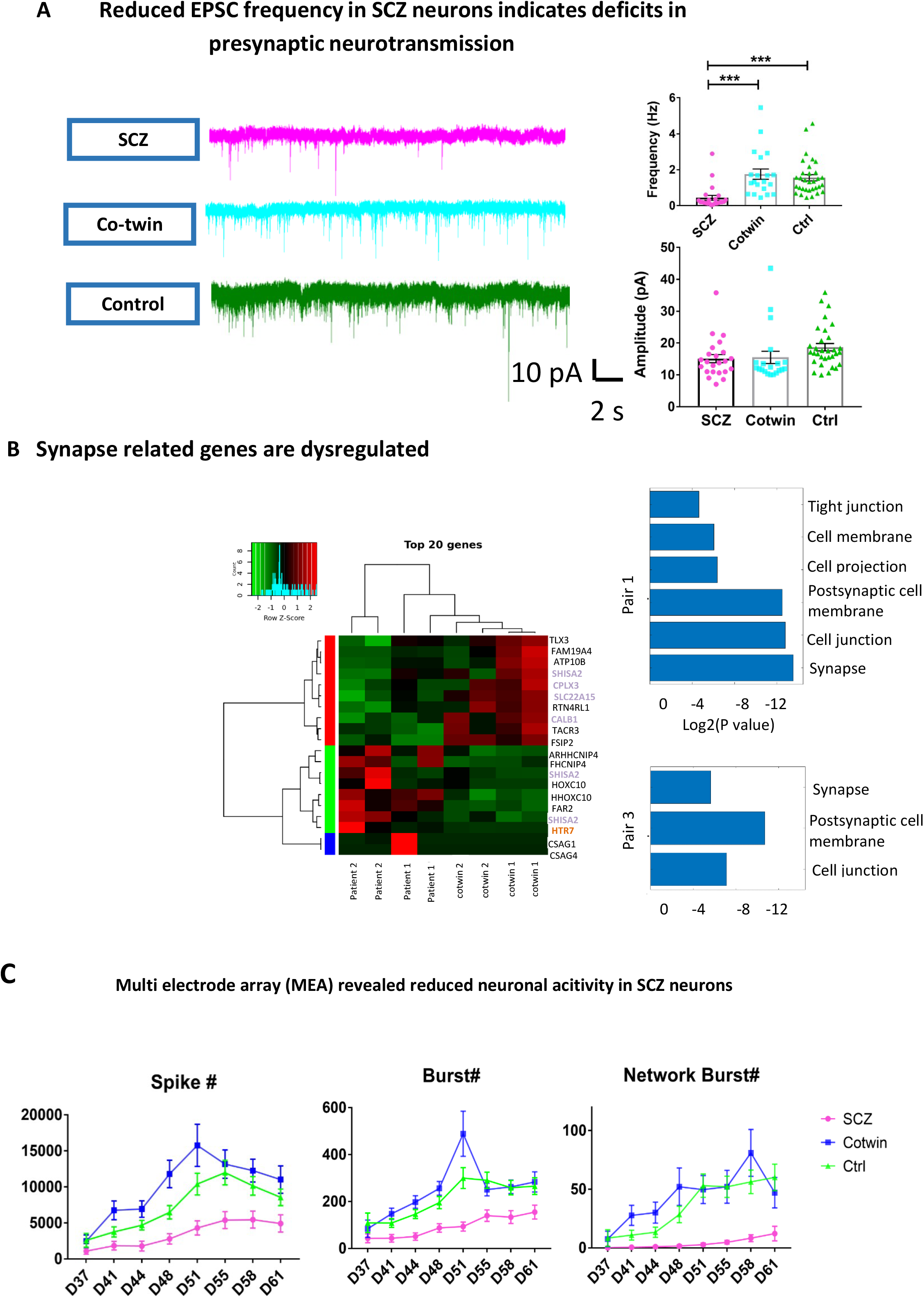
Synaptic deficiency in the affected twin siblings. A. A reduction in the rate of synaptic events measured in patch-clamp experiments was observed in the affected twin sibling but not in the unaffected co-twin sibling. On the left are representative traces and on the right, the averages are presented. The amplitude of the synaptic events was not affected, implying that the changes are mostly related to the pre-synapse. B. On the left are the top 20 dysregulated genes when pooling the affected twins (both siblings) and comparing them to the healthy twins. Many genes that are synapse-related are in the top 20 genes (these genes are marked in purple). In red is a gene that has been associated with schizophrenia before^45, 46^. On the right are the dysregulated pathways when comparing the neurons derived from the affected twin sibling to the unaffected twin sibling for both pairs of discordant twins. The top 3 dysregulated pathways in both pairs are “Synapse”, ”Postsynaptic cell membrane”, and “Cell junction,” despite there being few shared dysregulated genes between the 2 pairs of discordant twins. C. Multi-electrode array experiments show that the number of spikes and the number of bursts are reduced in the neurons derived from the schizophrenia patients but increased in the unaffected co-twin siblings. This may contribute to the mechanisms by which these siblings are not affected by schizophrenia. *** represents p<0.001.

To elucidate whether the pre-synaptic deficit is related to the neuronal activity, we next performed recordings using a multi-electrode array. The neurons derived from the affected twins with schizophrenia had a reduced number of spikes, a reduced number of bursts, and a reduced number of network bursts throughout the differentiation period). Interestingly, the neurons derived from the unaffected co-twin exhibited an increase in the number of spikes and the number of bursts. This increased excitability may act as a protective mechanism for a less connected neuronal network that exhibits more spontaneous activity and compensates for the reduced connections.

### Intrinsic hypoexcitability of schizophrenia patient-derived neurons

To explore whether the reduced neural activity of schizophrenia neurons is rooted in an intrinsic deficit, we next measured the number of action potentials that were evoked in 20 voltage depolarization steps (see Methods section). The number of action potentials that were evoked in the affected twins was significantly reduced compared to the controls and the unaffected co-twins. Additionally, the neurons derived from the unaffected twins were in an intermediate state and they too were significantly hypoexcitable compared to the control neurons (Fig. 5A left for representative traces and Figure 5A right for averages over 38 control neurons, 30 neurons derived from the unaffected co-twins, and 35 derived from the affected twin). Analysis of the spike shape also revealed 3 different groups with distinct states: the neurons derived from the healthy controls, the neurons derived from the unaffected co-twins that were in an intermediate state between the affected twin and the healthy controls, and the neurons derived from the affected twin siblings. Figure 5B shows that the width of the action potential was wider in the neurons derived from the affected twins and also wider, but to a smaller extent, in the unaffected co-twin when compared to the control neurons. The spike amplitude was smaller in the neurons derived from the affected twin compared to the other 2 groups. The threshold for evoking an action potential was unaffected. And finally, the amplitude of the fast AHP was smaller in neurons derived from both the affected and unaffected twins compared to the neurons derived from the healthy control neurons.

**Figure 5.**
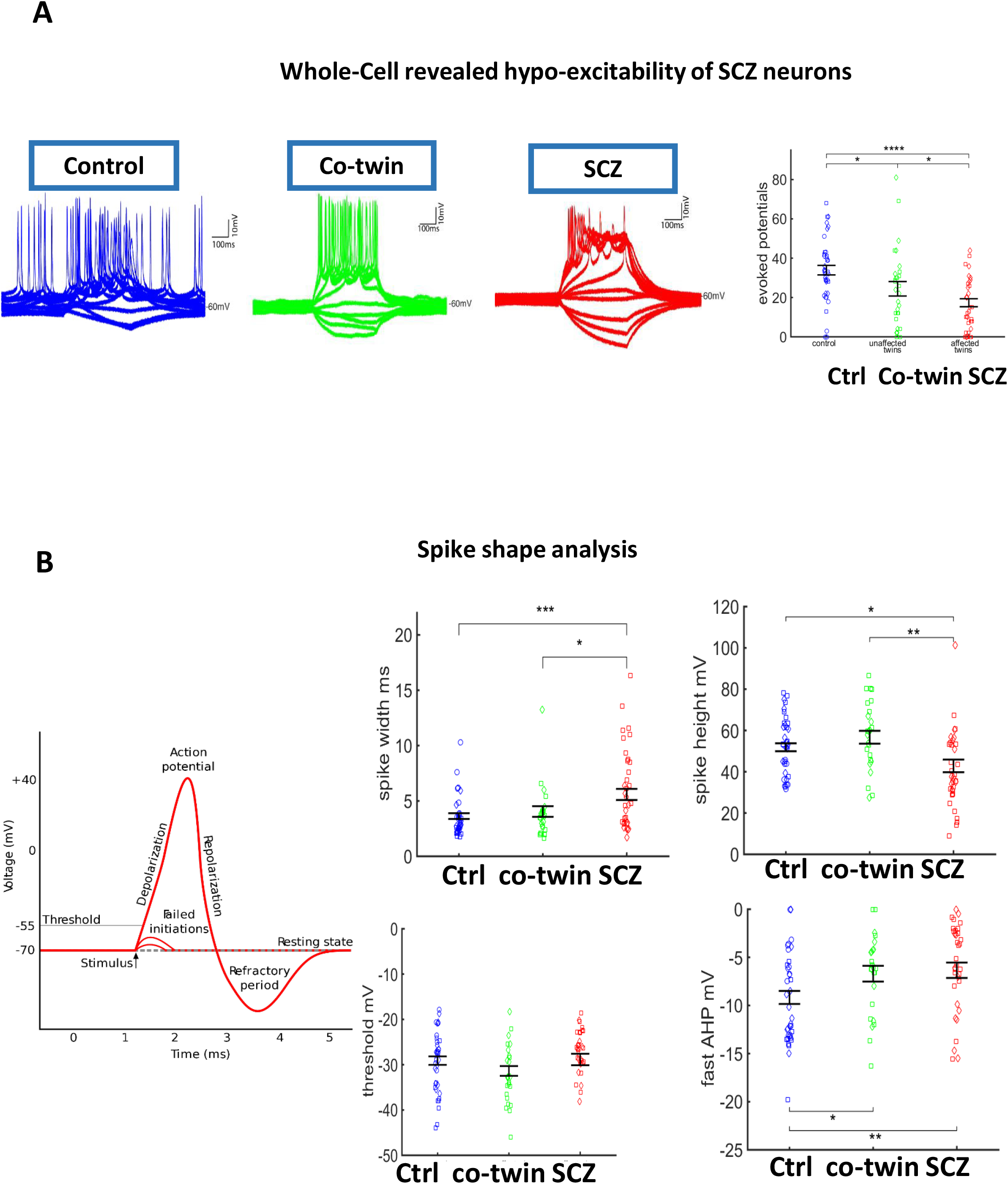
Hypoexcitability of neurons derived from schizophrenia patients. A. The neurons derived from the schizophrenia patients produced fewer evoked action potentials than the neurons derived from the other 2 groups. The neurons derived from the unaffected twin siblings produced more evoked action potentials than those from the affected twin siblings but fewer than the neurons derived from the healthy twin pairs. On the left are representative trace recordings. On the right, the averages of the total evoked potentials (see Methods) are presented. B. Spike shape analysis. The spike width was wider in the neurons derived from the affected twin siblings compared to the other 2 groups. The spike amplitude was smaller in the neurons derived from the affected twin siblings compared to the neurons derived from the other 2 groups. There was no significant change in the threshold for eliciting an action potential between the 3 groups. The amplitude of the fast afterhyperpolarization (AHP) was smaller in the neurons derived from both the affected twin siblings and the unaffected twin siblings. * represents p<0.05, **** represents p<0.0001. *represents p<0.05, ** represents p<0.01, *** represents p<0.001, **** represents p<0.0001.

### Three distinct groups were observed throughout the maturation of the DG neurons

We next asked whether the maturation delay was exhibited in earlier time points. We analyzed the recordings at an earlier time point of 4 weeks’ post-differentiation. The neurons were immature at this stage and most of them still did not have much evoked or spontaneous activity. However, measuring some neurophysiological properties we could see that the 3 groups had three different rates of maturation. At 4 weeks, the neurons derived from the affected twins had a smaller capacitance than the other 2 groups (Fig. 6A, p=4.6e-7 compared to neurons derived from the unaffected co-twins, p=3.8e-13 compared to neurons derived from the healthy twins, and p=0.19 between neurons derived from the control and the unaffected twins), but at 2 months 3 distinct groups were significantly different from each other (Fig. 6A, p=0.0029 for neurons derived from the schizophrenia patients compared to the unaffected co-twins, p=8e-15 for neurons derived from the affected twins compared to the healthy twins, and p=1.3e-9 for neurons derived from the unaffected co-twin compared to the healthy twins). The input resistance at 4 weeks was significantly and extensively larger in the neurons derived from the affected twins (Fig. 6B, p=4.5e-5 for neurons derived from the schizophrenia patients compared to the unaffected co-twins, p=2.1e-10 for neurons derived from the affected twins to the healthy twins, and p=0.04 for neurons derived from the unaffected co-twin compared to the healthy twins), but at 2 months the neurons derived from the affected and non-affected co-twins were significantly larger than the neurons derived from the controls (Fig. 6B, p=0.26 for neurons derived from the schizophrenia patients compared to the unaffected co-twins, p=3.7e-9 for neurons derived from the affected twins compared to the healthy twins, and p=2.8e-5 for neurons derived from the unaffected co-twins compared to the healthy twins). Also, notably, the input resistance drops for the schizophrenia twins by more than two-fold. This suggests together with the other data in this figure, suggests that early on the changes are even stronger and that schizophrenia is in fact a neurodevelopmental disorder although the onset is mainly during early adulthood.

**Figure 6.**
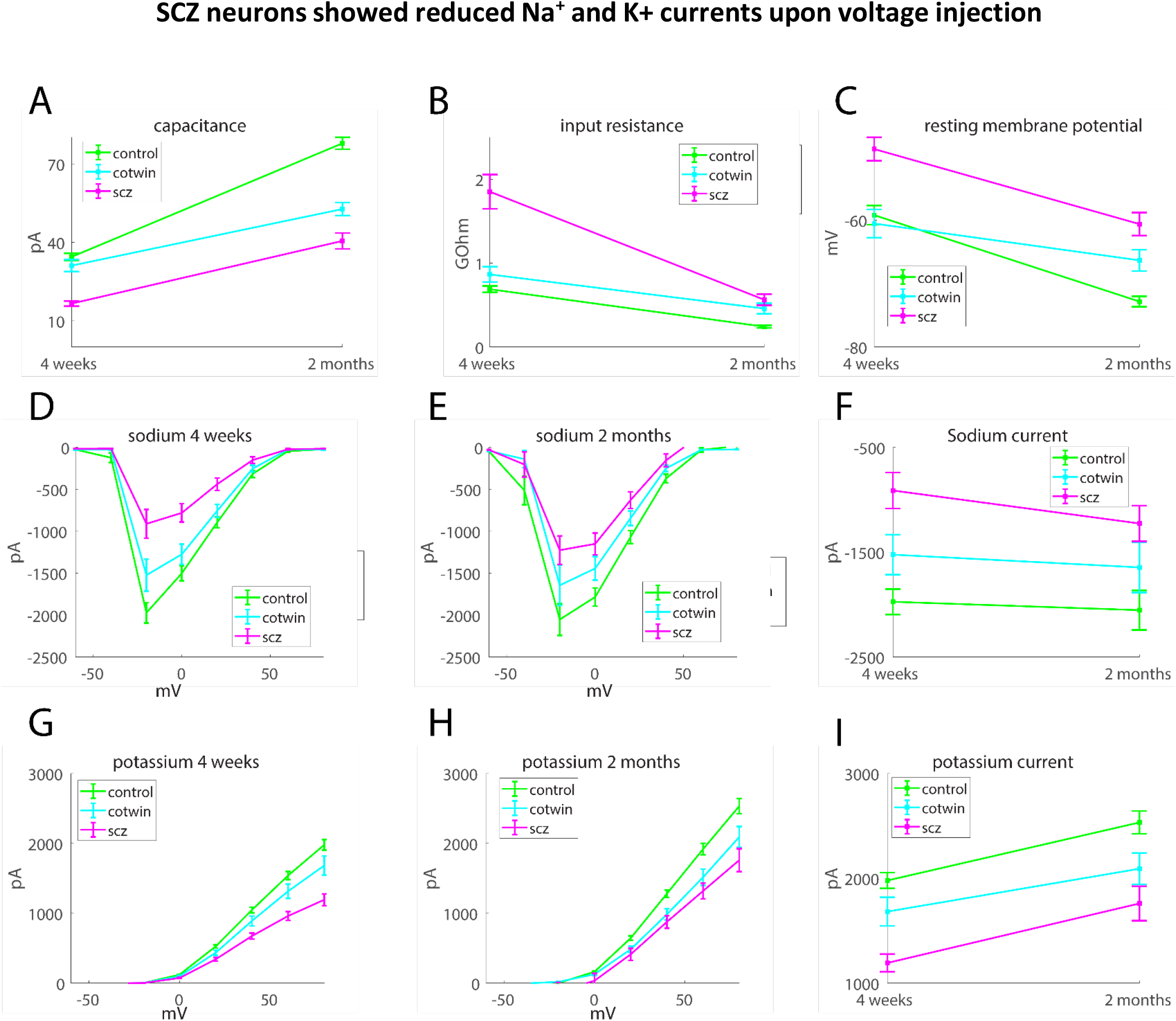
Three groups were observed when analyzing neurophysiological features throughout the maturation of the DG granule neurons. A. The capacitance measurements of the 3 groups (neurons derived from the affected twin siblings, neurons derived from the unaffected co-twin siblings, and neurons derived from the healthy twins) at 4 weeks and 2 months. B. The input resistance measurements of the 3 groups at 4 weeks and 2 months. C. The resting membrane potential of the 3 groups at 4 weeks and 2 months. D. The sodium currents of the 3 groups at 4 weeks. E. The sodium currents of the 3 groups at 2 months. F. The sodium currents at -20 mV of the 3 groups at 4 weeks and 2 months. G. The potassium currents of the 3 groups at 4 weeks. H. The potassium currents of the 3 groups at 2 months. F. The potassium currents at 20 mV of the 3 groups at 4 weeks and 2 months.

The resting membrane potential was much more depolarized in the neurons derived from the affected twins at 4 weeks (Fig. 6C p=1.7e-4 for neurons derived from the schizophrenia patients compared to the unaffected co-twins, p=6.1e-5 for neurons derived from the affected co-twins to the healthy twins, and p=0.62 for neurons derived from the unaffected co-twins compared to the healthy twins), and at 2 months the 3 groups were significantly different from each other, with the neurons derived from the affected twins having the most depolarized threshold, then the neurons derived from the unaffected twins, and finally the most hyperpolarized resting membrane potential belonging to the neurons derived from the healthy controls (Fig. 6C p=0.025 for neurons derived from the schizophrenia patients compared to the unaffected twins, p=4.5e-10 for neurons derived from the affected twins to the healthy twins, and p=1.6e-4 for neurons derived from the unaffected co-twins compared to the healthy twins).

Both the sodium and the potassium currents started as smaller currents in the neurons derived from the schizophrenia twins, with the unaffected co-twin in an intermediate state between the affected twins and the control (for sodium at -20 mV, p=4.7e-6 control-schizophrenia patients, p=0.024 co-twin-schizophrenia patients, p=0.045 co-twin-controls, for potassium at 20 mV, p=5.5e-6 control-schizophrenia, p=0.056 co-twin-schizophrenia, p=0.039 co-twin-control). At 2 months, the neurons derived from the schizophrenia patients still had reduced sodium currents (Fig. 6D-I, For sodium at -20 mV, p=0.0053 control-schizophrenia, p=0.24 co-twin-schizophrenia, p=0.158 co-twin-control, For potassium at 20 mV, p=0.0034 control-schizophrenia, p=0.5 co-twin-schizophrenia, p=0.005 co-twin-control). Overall, this analysis shows that the neurons derived from the affected twins had a very delayed maturation compared to the other 2 groups. At 2 months, when the neurons were more mature, most neurophysiological features split into 3 groups, where the neurons derived from the unaffected co-twins were in an intermediate state between the other two groups.

## Discussion

In this study, we differentiated DG granule hippocampal neurons from twin sets that were discordant for schizophrenia and compared them to control neurons that were derived from healthy twin sets. The neurons that were derived from the schizophrenia patients were smaller (less arborized with decreased capacitance), hypoexcitable (as measured in patch-clamp as less evoked action potentials and on MEAs as reduced spontaneous activity and reduced number of bursts), with immature features of the action potential. They had reduced sodium and potassium currents and a decrease in their synaptic activity. They had a more depolarized resting membrane potential and higher membrane resistance. The neurons derived from the twin siblings that were unaffected by schizophrenia were in an intermediate state between their affected siblings and the healthy controls in many of the neurophysiology aspects that we measured. For example, the neurons from the unaffected siblings were also less arborized with reduced capacitance when compared to the control neurons but more arborized and with a higher capacitance when compared to the neurons derived from their siblings who were suffering from schizophrenia. Their membrane resistance was smaller than their affected twin but larger than the control twins. Similarly, their resting membrane potential was in between the schizophrenia patients and the controls. Their excitability when measured by the total number of action potentials was reduced compared to the control neurons but higher than the neurons derived from their affected twin siblings. However, when measuring the spontaneous activity by MEAs, they were more excitable than the controls (and both were more excitable than the affected twins). Their sodium currents were also larger than their affected twin sibling but smaller than the currents of the control neurons. Their synaptic currents were unaffected compared to the control neurons.

The synaptic connectivity was measured by the rate of EPSCs and was severely reduced in the affected twins compared to the other 2 groups. When calculating the 20 most differentially expressed genes between the twin sets with schizophrenia compared to the control twin sets, 6 were synapse-related. One of the dysregulated pathways in this comparison was the “anterograde trans-synaptic signaling.” Furthermore, we compared the affected twins to the unaffected twins, each twin set separately. For pair number 1, the 3 most dysregulated pathways were “Synapse,” “Cell junction,” and “Postsynaptic cell membrane.” In pair 3, the most dysregulated functional annotations were similarly “Cell junction,” “Postsynaptic cell membrane,” and “Synapse.” Clearly, a synaptic impairment was the strongest phenotype of both the schizophrenia patients compared to their twin siblings both when measuring the synaptic signaling with patch-clamp and at the gene expression level. Synaptic impairment has been linked with schizophrenia in animal studies ^48^, in post mortem tissue^49, 50^, and in patient-derived neurons^51^. Here we confirm that there is also a synaptic impairment in our DG granule patient-derived neurons. Our results are probably the strongest indication that synaptic deficit is indeed a prominent phenotype of neurons derived from schizophrenia patients, as it is observed in the affected twins when measured in electrophysiology and not in the unaffected twins. However, at the gene expression level, there is a synaptic dysregulation also in the co-twins, but it is less prominent.

The neurons derived from the schizophrenia patients also exhibited a decreased number of evoked potentials, and the unaffected sibling was in an intermediate state between the affected twin and the healthy controls. A decreased number of evoked action potentials has also been previously reported in schizophrenia patients^33^. Interestingly, schizophrenia shares multiple genomic associations with bipolar disorder^52-54^, but DG granule neurons derived from bipolar disorder patients are hyperexcitable whereas those derived from schizophrenia patients are hypoexcitable^27, 55, 56^. Our study shows that the unaffected twin also had fewer evoked potentials with current injections but, on the other hand, it had more spontaneous activity when measured in MEAs. This increased spontaneous activity may help to rescue these individuals as it strengthens network activity in a network where the synapses are affected.

The RNA sequencing results indicate the DG development pathway is also dysregulated when comparing the control twins and the affected twins (both affected and unaffected twin siblings). Indeed, we see that the cells of the affected twins were smaller, with a reduced capacitance. The smaller membrane resistance observed in the affected twins also signifies that there is a delay in the expression of ion channels such as the inward rectifying potassium channels and this is further supported by the more depolarized resting membrane potential. A delay in the development of gray and white matter in adolescent patients with schizophrenia has also been previously reported ^57, 58^. Common genetic variants that affect rates of brain growth or atrophy, including the hippocampus, showed genetic overlap with schizophrenia in a recent meta-analysis of changes in brain morphology across the lifespan ^59^. The DG newly born granule neurons are thought to be extremely important to the DG circuitry ^60, 61^. Newly born DG granule neurons are formed throughout our lives, so this immature phenotype affects both the development and the integration of these delayed developing DG neurons continuously throughout the patients’ lives.

Our study demonstrates several phenotypes of DG granule neurons derived from schizophrenia patients and their unaffected twin siblings that demonstrate only a partial list of these phenotypes. The synaptic impairment is a noticeable phenotype that appears both in the electrophysiological recordings and in the transcriptional analysis. This phenotype is much less prominent in the unaffected twin, which further indicates the importance of hippocampal synaptic impairment in the mechanism of this disease.

That both the unaffected and affected twin pair are morphologically and electro-physiologically different from control twins suggests an underlying germline genetic defect that is further supported by transcriptional profile differences. Epigenetic modifications can also be responsible for the disease onset in one of the patients. However, the ability to detect a phenotype using iPSC-derived neurons implies that genetic differences between the patients play a role as well. This means that the phenotypical changes that we measured in the derived neurons, may show relevant mechanisms for the symptoms in the individuals. It is important to mention, that the unaffected co-twins were later diagnosed with a depressive disorder. Furthermore, that we find morphological and electro-physiological differences between discordant twin pairs, that are further confirmed by transcriptional differences suggests that in addition to the proposed germline changes between schizophrenia and neurotypical controls there are likely somatic changes in the genomes of the affected or unaffected pair that occurred after the twins split following fertilization, We are currently conducting whole genome sequencing on blood and fibroblasts to test the validity of this speculation.

## Acknowledgments

The authors would like to thank K.E. Diffenderfer for technical assistance and M.L. Gage for editorial comments. They would also like to acknowledge the Salk Institute Stem Cell Core, Waitt Biophotonics Core, and Next Generation Sequencing Core for technical support. The authors acknowledge the Bioimaging Unit, Faculty of natural sciences, University of Haifa. Funding to the cores provided in part by NIH-NCI CCSG: P30 014195. This work was supported by the National Cancer Institute (Grant No. P30 CA014195 [to FG]), the National Institutes of Health (Grant No. R01 AG05651 [to FG]), the National Cooperative Reprogrammed Cell Research Groups (Grant No. U19 MH106434 [to FG]), the JPB Foundation (to FG), Annette C. Merle-Smith (to FG), and the Robert and Mary Jane Engman Foundation (to FG). This work was supported by the Zuckerman STEM leadership program (to SS) and the Israeli Science Foundation (Grants No. 1994/2 and 3252/21 to SS).

